# NOX4 inhibition promotes the remodeling of dystrophic muscle

**DOI:** 10.1101/2022.01.08.475493

**Authors:** David W. Hammers

**Author notes:** Corresponding Author **Correspondence:** David W. Hammers, 1200 Newell Dr., ARB R5-234, Gainesville, FL 32610-0267.

## Abstract

The muscular dystrophies (MDs) are genetic muscle diseases that result in progressive muscle degeneration followed by the fibrotic replacement of affected muscles as regenerative processes fail. Therapeutics that specifically address the fibrosis and failed regeneration associated with MDs represent a major unmet clinical need for MD patients, particularly those with advanced stage disease progression. The current study investigates targeting NAD(P)H oxidase (NOX) 4 as a potential strategy to reduce fibrosis and promote regeneration in disease-burdened muscle that models Duchenne muscular dystrophy (DMD). NOX4 is elevated in the muscles of dystrophic mice and DMD patients, localizing primarily to interstitial cells located between muscle fibers. Genetic and pharmacological targeting of NOX4 significantly reduces fibrosis in dystrophic respiratory and limb muscles. Mechanistically, NOX4 targeting decreases the number of fibrosis-depositing cells (myofibroblasts) and restores the number of muscle-specific stem cells (satellite cells) to their physiological niche, thereby, rejuvenating muscle regeneration. Furthermore, acute inhibition of NOX4 is sufficient to induce apoptotic clearing of myofibroblasts within dystrophic muscle. These data indicate that targeting NOX4 is an effective strategy to promote the beneficial remodeling of disease-burdened muscle representative of DMD and, potentially, other MDs and muscle pathologies.

**SIGNIFICANCE STATEMENT:** Muscular dystrophies are progressive muscle diseases. Therapeutics capable of combating the severe fibrosis that replaces functional muscle in these devastating diseases is a major unmet clinical need, particularly for the treatment of older patients. The current work reveals that targeting NOX4 in dystrophic muscle promotes the beneficial remodeling of disease-burdened musculature. This is achieved by the removal of disease-causing cells, known as myofibroblasts, which results in reduced muscle fibrosis and rejuvenation of muscle regeneration. NOX4-targeting strategies, therefore, represent remodeling therapeutics capable of improving muscle disease caused by muscular dystrophy, and, likely, other muscle pathologies.

## INTRODUCTION

Muscular dystrophies (MDs) are genetic muscle diseases characterized by progressive skeletal muscle degeneration and replacement of functional musculature with an aberrant fibrotic extracellular matrix (ECM). Duchenne MD (DMD), the most prevalent of MDs, is a fatal, childhood-onset X-linked disease that affects ∼1:5000 males (1). DMD is caused by mutations in the *DMD* gene resulting in complete loss of dystrophin (2), a protein that stabilizes the sarcolemma during contractile activity by providing a link between the muscle cytoskeleton and surrounding ECM (3). Patients of this devastating disease typically progress to loss of ambulation within the second decade of life and death by the age of 30. During the course of disease progression, DMD patients exhibit a profound expansion of a fibrotic and fatty ECM as muscle fibers are lost (4). Therapeutics capable of slowing or reversing this ECM expansion are a major unmet clinical need, particularly for disease management of older DMD patients.

The identification of treatments that effectively improve dystrophic muscle has been largely hindered by limited knowledge of the cellular mechanisms responsible for the development of muscle fibrosis and impairment of muscle regeneration associated with the disease. Under normal conditions, skeletal muscle displays robust regeneration following injury, which is largely mediated by muscle-resident stem cells, known as satellite cells (5, 6). The process of muscle regeneration also requires an array of chemical and physical cues provided by immune cells, myogenic cells, fibroblasts, and other cellular populations (7–9). These factors are temporally orchestrated to form new muscle that is accommodated with adequate vasculature, innervation, and ECM structural support, leading to resolution of the injury response and return to homeostasis (5, 7, 10). Indeed, perturbations to any of these components contributes to maladaptive muscle regeneration (7, 9, 11, 12). In dystrophic muscle, the continuous and asynchronous combination of degeneration and regenerative processes within the same muscle disrupts the timing of these events (13). This ultimately leads to regenerative impairments and progressing fibrosis that characterize the disease burden of late-stage dystrophic muscle.

Upregulation of NAD(P)H oxidase (NOX) 4 was identified in diseased muscle of D2.*mdx* mice, a severe mouse model of DMD (14–16), using a previously published transcriptomic data set (17). NOX4 is a reactive oxygen species (ROS)-generating enzyme that has been implicated in fibrosis of the lung, kidney, liver, and heart [reviewed in (18)]. While elevated *Nox4* expression has been noted in dystrophic mouse hearts (19), NOX4 has not been previously studied in regards to skeletal muscle fibrosis. The current study sought to determine if the targeting NOX4 is an effective strategy to prevent fibrosis and enhance regeneration in dystrophic muscle. Herein, it is shown that NOX4 localizes primarily to interstitial cells of dystrophic muscle that resemble active ECM-secreting cells, known as myofibroblasts, which are typically only found in damaged or diseased tissue (20). The targeting of NOX4 both by genetic ablation and pharmacological inhibition promotes the beneficial remodeling of diseased muscle by reducing muscle fibrosis. Importantly, NOX4 targeting substantially reduces myofibroblasts within disease-burdened muscle, restores the localization of satellite cells to their physiological niche, and increases evidence of muscle regeneration. These data implicate NOX4 in the development of MD-associated skeletal muscle pathology, and demonstrate that targeting NOX4 is an effective strategy to promote beneficial remodeling of dystrophic muscle.

## RESULTS

### NOX4 is an anti-fibrotic target in in D2.mdx skeletal muscle

The heightened skeletal muscle fibrosis of D2.*mdx* mice makes this emerging mouse model of DMD better suited to investigate mechanisms contributing to muscle fibrosis than *mdx* mice on C57-based genetic backgrounds (14). To identify potential gene targets that may be exploited as anti-fibrotic therapies, a previously published transcriptomic dataset (17) was queried for ECM/fibrosis-associated genes that are significantly upregulated in D2.*mdx* quadriceps muscle, relative to wild-type DBA/2J (D2.WT) values (p < 0.05). Of 26 genes identified (**Figure 1A**), the majority consisted of ECM components, including 10 isoforms of collagen, 4 members of matrix metallopeptidases (MMPs; a class of matrix remodeling enzymes), the collagen crosslinking enzymes *Lox* and *Loxl1*, and other ECM-related genes, including *Fn1* (fibronectin), *Ltbp2, Postn* (periostin), and *Spp1* (osteopontin). Identified fibroblast-lineage markers include *Pdgfra* [platelet-derived growth factor receptor (PDGFR) α], *Ctgf, Vim* (vimentin), *S100a4*, and *Acta2* [α-smooth muscle actin (SMA)].

**Figure 1.**
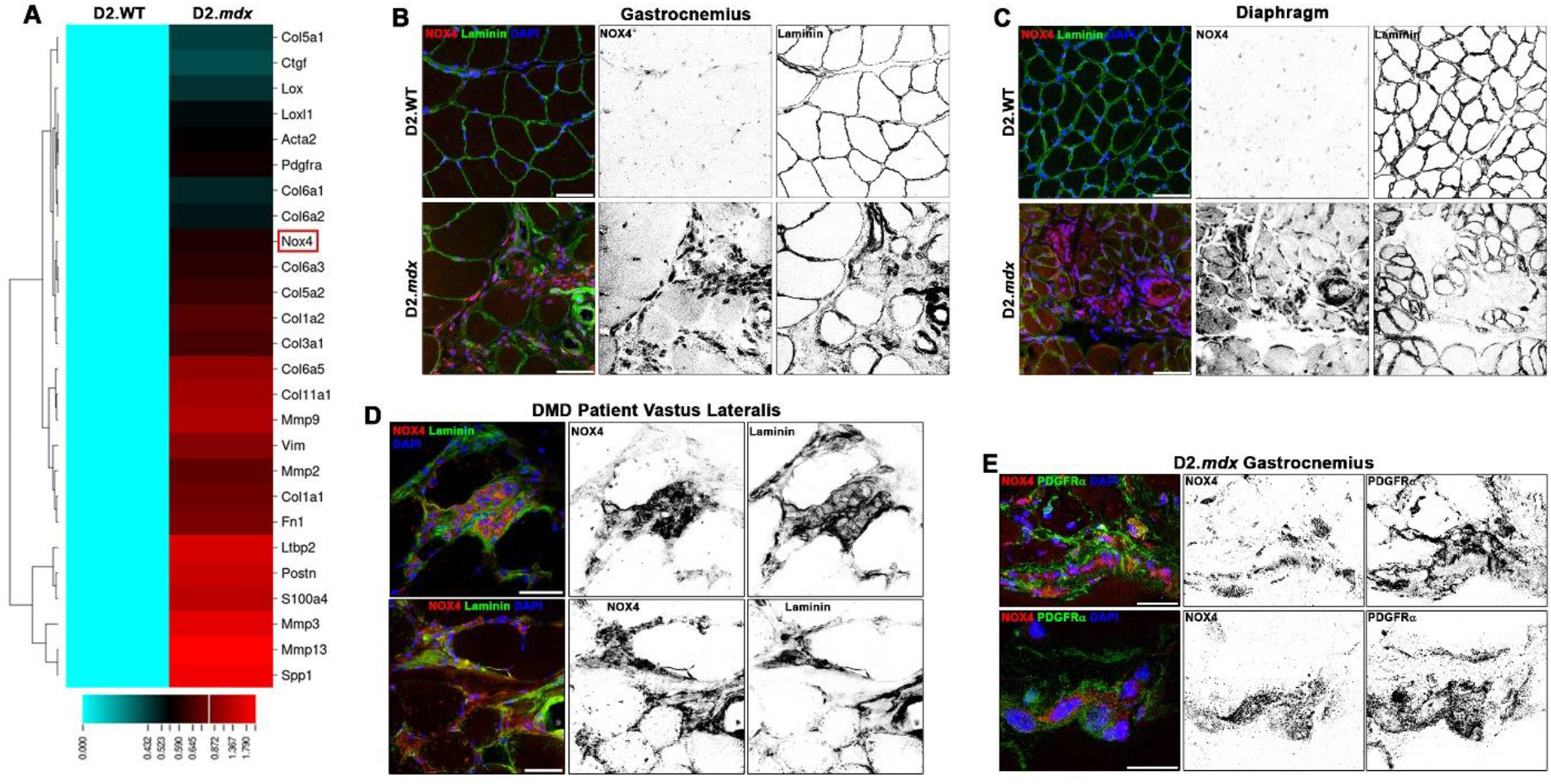
NOX4 is increased in dystrophic skeletal muscle. (**A**) Fibrosis-associated genes differentially expressed in 4 month-old male wild-type DBA/2J (D2.WT) and dystrophic D2.*mdx* quadriceps muscles were queried from transcriptomic data [p < 0.05; depicted as log(fold-change); published in ref. (17)]. *Nox4* is highlighted by the red box. Immunofluorescent detection of NOX4 in the (**B**) gastrocnemius and (**C**) diaphragm muscles of D2.WT and D2.*mdx* mice, as well as (**D**) the vastus lateralis of DMD patients, reveals NOX4 is primarily localized to interstitial cells of dystrophic muscle. (**E**) NOX4 is localized to fibroblast-lineage cells, labeled by PDGFRα, in the dystrophic muscle of D2.*mdx* mice. Scale bars represent (**B-C**) 100 or (**D-E**) 50 µm.

An interesting find amongst this query was the increased expression of *Nox4*, which encodes the intracellular ROS-generating enzyme, NOX4. NOX4 has been identified as an anti-fibrosis target in several tissue types (21–28), but has not previously been associated with skeletal muscle fibrosis. Importantly, safe clinical-stage small-molecule inhibitors of this enzyme have been developed (29). Immunoblotting confirms NOX4 protein is elevated in D2.*mdx* skeletal muscle (**Figure S1A**), which is localized at high concentrations in the interstitial regions between muscle fibers of D2.*mdx* mice, as shown in the gastrocnemius (**Figure 1B**) and diaphragm (**Figure 1C**). This NOX4 staining pattern is also found in the muscles of DMD patients (**Figure 1D**). These NOX4-expressing cells observed in D2.*mdx* muscle also stain positive for PDGFRα (**Figure 1E**), a marker of fibroblast-lineage cells, including fibro-adipogenic progenitors (FAPs), fibroblasts, and myofibroblasts (30–33). Many of the PGDFRα^+^ cells found in NOX4-enriched interstitial regions also express vimentin and SMA (**Figure S1B**), markers of myofibroblasts (34). NOX4-expressing PDGFRα^+^ cells are also found in the muscles of mice that model limb-girdle MD (LGMD) 2B (dysferlinopathy; **Figure S1C**) and 2F (δ-sarcoglycan deficiency; **Figure S1D**), therefore these observations are not restricted to dystrophin-deficient muscle. This evidence suggests the hypothesis that NOX4 is potentially involved in the development of muscle fibrosis in the MDs.

The role of NOX4 in the development of muscle fibrosis was directly tested by the generation of a *Nox4* knockout mouse line on the D2.*mdx* background (Nox4^KO^:*mdx*). Wild-type littermates of this line (Nox4^WT^:*mdx*) are indistinguishable from mice of the D2.*mdx* colony and exhibit NOX4 staining in the muscle interstitium, whereas Nox4^KO^:*mdx* mice show no NOX4 immunoreactivity (**Figure S2A**). At 3 months of age, when D2.*mdx* mice begin to show fibrosis during recovery from widespread muscle degeneration (14), Nox4^WT^:*mdx* and Nox4^KO^:*mdx* littermates show no difference in muscle fibrosis, as assessed by histological evaluation using picrosirius red staining of the diaphragm and gastrocnemius (**Figure S2B–D**). Therefore, NOX4 does not appear to affect ECM production associated with tissue repair from injury. At 6 months of age, when D2.*mdx* muscle displays robust fibrotic progression independently of reparative activities (14), Nox4^KO^:*mdx* diaphragms and gastrocnemius muscles exhibit significantly less fibrosis than those of the Nox4^WT^:*mdx* littermates (**Figure 2A-C**). While Nox4^WT^:*mdx* muscles show the expected increase in fibrosis from 3 months to 6 months, Nox4^KO^:*mdx* muscles actually reduce muscle fibrosis during this timeframe. Thus, loss of NOX4 prevents the progressive accumulation of fibrosis in D2.*mdx* skeletal muscle, possibly by facilitating resolution of the post-injury response. NOX4-ablated diaphragms and extensor digitorum longus (EDL) muscles also exhibit significantly increased muscle function over those of Nox4^WT^:*mdx* mice (**Figure 2D-E**), demonstrating that functional muscle is better maintained as fibrosis is reduced. Furthermore, pharmacological targeting of NOX4 phenocopies this anti-fibrotic effect of NOX4 ablation, as 3 months of treatment with the NOX1/4 inhibitor GKT831 (21, 29, 35) also reduces D2.*mdx* muscle fibrosis, compared to vehicle treatments, when evaluated at 6 months of age (**Figure 2A-C,F**). Collectively, these data indicate that targeting NOX4 is an efficacious means to control the fibrotic replacement of dystrophic muscle.

**Figure 2.**
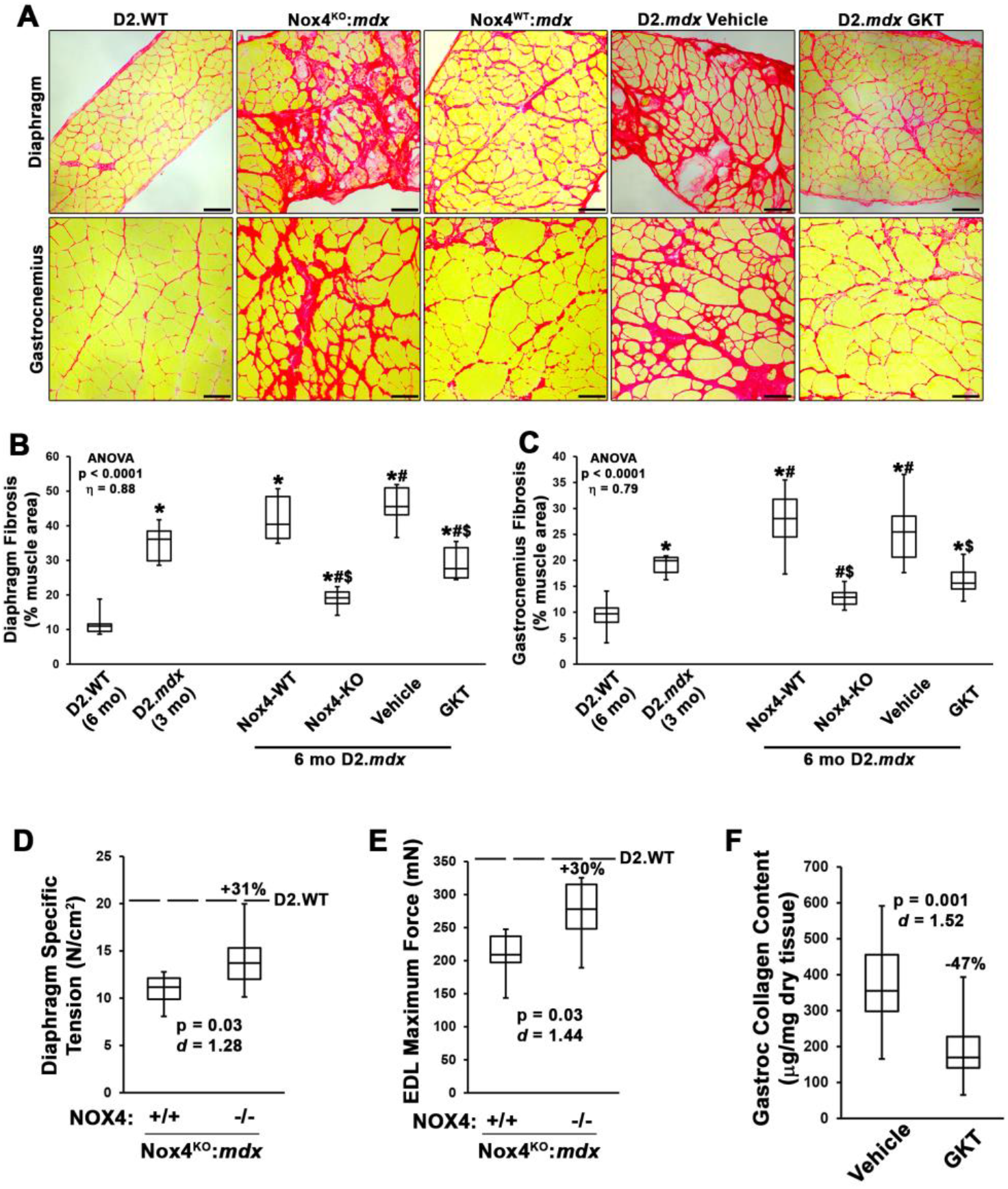
Targeting NOX4 reduces fibrosis in dystrophic muscle. (**A**) Representative picrosirius red staining and image quantification of (**B**) diaphragm and (**C**) gastrocnemius muscles from 6 month-old D2.WT (n = 6), NOX4 wild-type (NOX4-WT) and knockout (NOX4-KO) littermates from a NOX4 KO line generated on the D2.*mdx* background (Nox4^KO^:*mdx*; n = 12), and D2.*mdx* mice that received vehicle or GK831 (GKT) treatments beginning at 3 months of age (n = 12-14). Fibrosis quantifications for 3 month-old D2.*mdx* mice are included to show baseline values for GKT treatment groups. Muscle function was assessed for the (**D**) diaphragm and (**E**) extensor digitorum longus (EDL) muscles of NOX4-WT (+/+) and NOX4-KO (-/-) littermates of the Nox4^KO^:*mdx* mouse line. (**F**) Biochemical quantification of collagen content was performed on gastrocnemius muscles of vehicle and GKT-treated D2.*mdx* mice. Data are presented as box-and-whisker plots with error bars representing minimum and maximum values. (**D-F**) Percent values indicate the difference between mean values of the two groups. Data were analyzed using (**B-C**) ANOVA followed by Tukey post-hoc tests (α = 0.05; effect size is reported as η^2^) or (**D-F**) unpaired, two-tailed Welch’s T-tests (α = 0.05; effect size is reported as Cohen’s *d*); *p < 0.05 vs. D2.WT values; #p < 0.05 vs. 3 mo D2.*mdx* values; $p < 0.05 vs. respective control group values.

### NOX4 targeting promotes dystrophic muscle remodeling through clearance of myofibroblasts

Myofibroblasts actively secrete ECM in tissues during repair and disease (20), and NOX4 is expressed by myofibroblasts in fibrotic lungs (24) and dystrophic muscle (**Figure 1C, S1D-E**). Therefore, it is likely that NOX4 targeting exerts its anti-fibrotic effects primarily by targeting myofibroblasts. In agreement, myofibroblast numbers, as determined by labeling with PDGFRα, vimentin, and SMA, are significantly reduced by both NOX4 ablation and GKT831 treatment (**Figure 3A**). Additionally, these NOX4-targeting strategies reduce the amount of periostin, an ECM component that is specifically expressed and secreted by myofibroblasts in muscle tissues (36), in D2.*mdx* muscle (**Figure 3B**). These data indicate that the anti-fibrotic efficacy of NOX4 inhibition in dystrophic muscle is due to the clearance of myofibroblasts, thereby, removing the source of pathological ECM deposition from the muscle.

**Figure 3.**
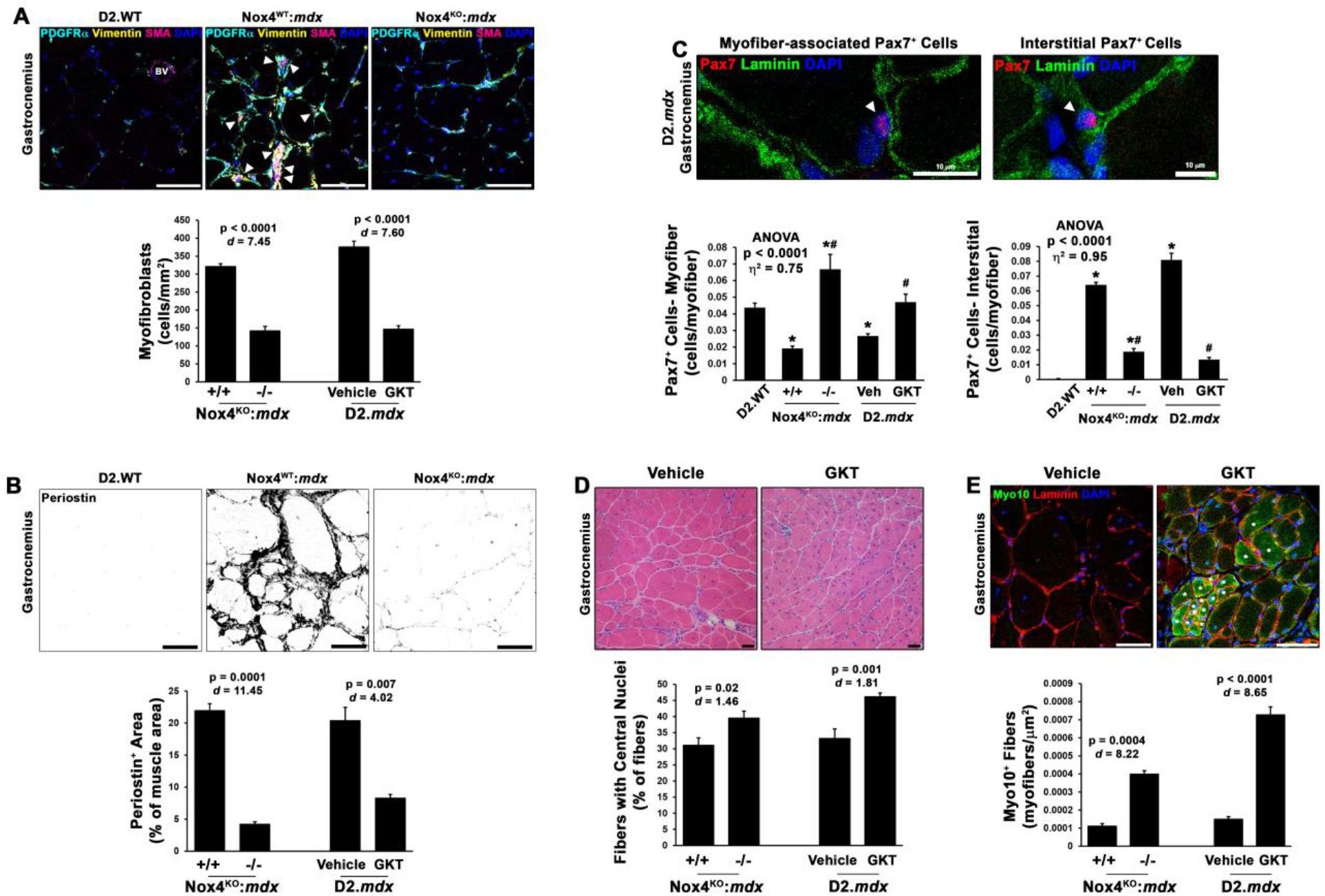
NOX4 targeting reduces myofibroblasts and increases regeneration in dystrophic muscle. Gastrocnemius muscles from 6 month-old Nox4^WT^:*mdx* (+/+), Nox4^KO^:*mdx* (-/-), and D2.*mdx* mice receiving vehicle (Veh) or GKT831 (GKT) treatments were assessed for indices of myofibroblast numbers and muscle regeneration (n = 6-12). Representative immunofluorescent images and quantifications are shown for (**A**) myofibroblast prevalence, as determined by co-labeling with PDGFRα, vimentin, and SMA (arrows mark myofibroblasts; BV = blood vessel), and (**B**) muscle periostin content. Pax7 immunofluorescence was performed to assess satellite cell numbers. Populations of myofiber-associated and interstitial Pax7^+^ cells were identified and affected by NOX4 targeting, as displayed by (**C**) representative images and population quantifications (arrows mark Pax7^+^ cells). Evidence of restarted muscle regeneration by NOX4 targeting was also found, as shown by number of fibers exhibiting (**D**) centrally-located nuclei and (**E**) staining for myosin X (Myo10; stars indicate fibers marked as recently regenerated). Data are displayed as mean±SEM and were analyzed using (**A-B**,**D-E**) unpaired, two-tailed Welch’s T-tests (α = 0.05; effect size is reported as Cohen’s *d*) or ANOVA followed by Tukey post-hoc tests (α = 0.05; effect size is reported as η^2^). *p < 0.05 vs. D2.WT values; #p < 0.05 vs. respective control group values. Unless otherwise indicated, scale bars indicate 50 µm.

Evidence suggests that myofibroblasts may inhibit muscle regeneration (37–39). In agreement with a suppressed regenerative capacity in disease-burdened D2.*mdx* muscle, Nox4^WT^:*mdx* and vehicle-treated gastrocnemius muscles exhibit significantly reduced numbers of Pax7^+^ satellite cells localized to their physiological niche between the sarcolemma and basal lamina of muscle fibers (**Figure 3C**). However, a substantial number of Pax7^+^ cells are found to be displaced into the dystrophic muscle interstitium (**Figure 3C)**, similar to what has been previously shown in DMD patient muscle (40) and murine cancer cachexia (41). In agreement with the beneficial remodeling of the dystrophic musculature by NOX4 targeting, NOX4 ablation and GKT831 treatments largely re-localize Pax7^+^ cells back to the periphery of muscle fibers (**Figure 3C**). Furthermore, this return of Pax7^+^ cells to their physiological niche coincides with increases in muscle fibers having central nuclei (**Figure 3D**) and expression of myosin X (**Figure 3E**), both indicators of recent muscle regeneration (10, 42). Therefore, the beneficial remodeling of dystrophic muscle by NOX4 inhibition also entails restoration of muscle regenerative capacity. These data depict a model whereby NOX4 targeting promotes the remodeling of dystrophic muscle via the clearance of myofibroblasts and alleviation of regeneration-inhibiting actions on satellite cells. Because the presumed mechanism attributed to myofibroblast clearance is by promoting their apoptosis (20), a short-term study was performed to investigate acute changes in muscle that result from NOX4 inhibition. Six month-old D2.*mdx* mice, whose muscles exhibit substantial disease burden (14), received vehicle or GKT831 treatments for seven days. Following this dosing period, the gastrocnemius muscles of GKT831-treated mice exhibited significantly more PDGFRα^+^ cells showing positive staining for active Caspase-3 (**Figure 4A**), an indicator of apoptosis. This demonstrates that the myofibroblast clearance achieved by NOX4 inhibition occurs, at least in part, via apoptosis. In agreement, muscles of GKT831-treated mice also exhibit decreased expression of the fibrosis/myofibroblast markers *Col1a2* and *Acta2*, as well as the pro-fibrotic and anti-regeneration markers of *Il6* and *Tgfb1*, compared to those of vehicle-treated mice (**Figure 4B**). These data reinforce the notion that NOX4 inhibition promotes the remodeling of dystrophic muscle by inducing myofibroblast apoptosis.

**Figure 4.**
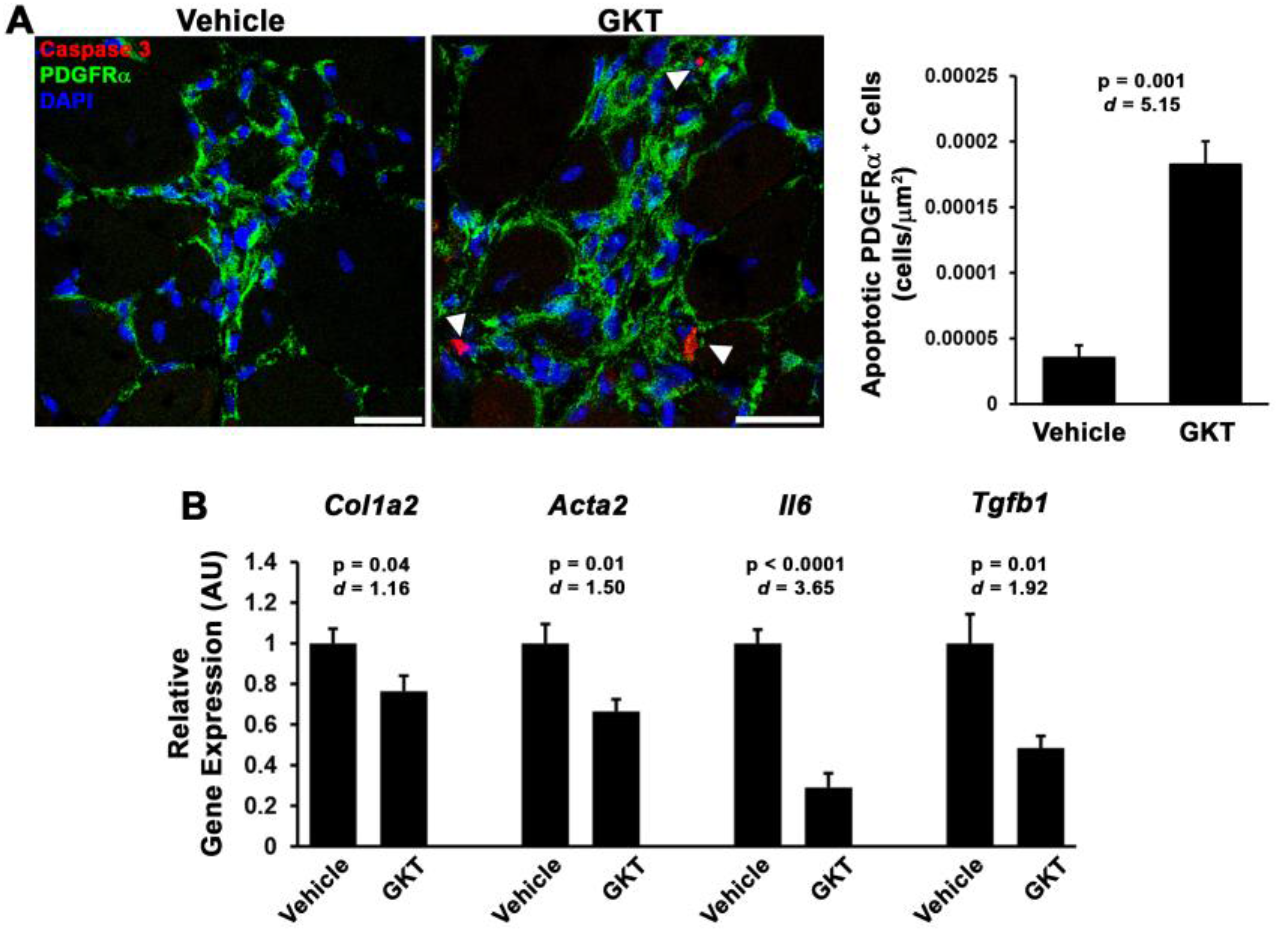
Short-term NOX4 inhibition induces myofibroblast apoptosis. Six month-old male D2.*mdx* mice were treated with vehicle or 60 mg/kg GKT831 (GKT) for seven days (n = 6). (**A**) Apoptosis of PDGFRα^+^ cells in vehicle and GKT-treated gastrocnemius muscles was assessed by immunofluorescent detection of active caspase-3. (**B**) Gene expression for *Col1a2, Acta2, Il6*, and *Tgfb1* was also measured in gastrocnemius muscles using real-time PCR. Data are displayed as mean±SEM and were analyzed using unpaired, two-tailed Welch’s T-tests (α = 0.05; effect size is reported as Cohen’s *d*).

## DISCUSSION

Therapeutics capable of remodeling the disease-burdened muscle of MD patients is a major unmet clinical need. This is particularly important for the treatment of older DMD patients who have undergone substantial replacement of muscle with pathological ECM. The current work identifies NOX4 as an efficacious target to promote the remodeling of dystrophic muscle by preventing fibrosis and enhancing regeneration. This was achieved using genetic ablation and, importantly, pharmacological inhibition with a clinical-stage drug. The efficacious effect of NOX4 targeting is primarily associated with a reduction of myofibroblasts in dystrophic muscle. This clearance of myofibroblasts appears to alleviate both the pro-fibrotic and anti-regenerative environment associated with disease-burdened muscle. These findings indicate that NOX4-targeting interventions represent potential remodeling therapeutics capable of improving the muscle disease state in DMD and other muscle diseases.

Myofibroblasts are specialized ECM-depositing cells that are activated to facilitate tissue repair; however, they contribute to tissue pathology in chronic diseases by producing excessive ECM, thereby, causing progressive fibrosis (20). A major finding of the current work is that the beneficial remodeling incurred by NOX4 targeting involves the clearance of myofibroblasts within the muscles. While similar results have been shown in lung fibrosis (24, 43), this is the first report, to my knowledge, of a treatment that primarily targets myofibroblasts in dystrophic muscle. The clearing of myofibroblasts by targeting NOX4 is a two-hit approach to promote beneficial remodeling of dystrophic muscle, as **a**) the cellular source of fibrosis is removed from the system, and **b**) negative regulation of regenerative efforts is alleviated, thereby allowing a stalled myogenic program to proceed.

The inhibitory effect of myofibroblasts on muscle regeneration may be mediated by multiple mechanisms. For instance, periostin, a myofibroblast-specific protein that is secreted during ECM production (36), appears to directly inhibit muscle regeneration (37), and its ablation enhances regeneration (18, 38). Similarly, NOX4 inhibition drastically reduces periostin content in dystrophic muscle (**Figure 3B**), which coincides with evidence of rejuvenated regeneration (**Figure 3D-E**). Furthermore, myofibroblasts may also release signaling molecules that impede satellite cell function, such as TGFβ (44, 45). In this study, many Pax7^+^ cells were found to be mislocalized to the interstitium of dystrophic muscle (**Figure 3C**), which were largely restored to a myofiber-associated localization upon the removal of myofibroblasts by NOX4 targeting. This suggests that myofibroblasts, whether by secreted factors or physical interactions, essentially halt the activity of satellite cells within dystrophic muscle, thus, contributing to the regenerative decline associated with DMD. Importantly, this arrest of satellite cell activity appears to be reversible, as was previously shown in cancer cachexia (41), which allows active regeneration of dystrophic muscle to be restarted upon alleviation of these inhibitory signals.

Currently, the landscape of DMD therapeutics is at a pivotal crossroad. Advancements in gene therapy have led to clinical trials evaluating the delivery of miniaturized dystrophin transgenes to the muscles of DMD patients (46). If successful, these therapies will transform DMD into a slower progressing disease that still exhibits progressive replacement of muscle with ECM, which is seen in Becker muscular dystrophy patients (47, 48). Furthermore, it is not clear how pre-existing disease burden will affect the delivery or efficacy of these gene therapies, particularly in the case of treating older DMD patients. Thus, therapeutics capable of remodeling disease-burdened muscle will remain a major clinical need for the management of DMD, whether as a monotherapies or in combination with gene therapy. The current study indicates that NOX4-inhibiting strategies represent effective remodeling therapeutics capable of reducing fibrosis and enhancing muscle regeneration in dystrophic muscle through the targeting of myofibroblasts. These findings will likely be broadly impactful for the treatment of both genetic and non-genetic muscle diseases that exhibit progressive fibrosis and failed regeneration.

## EXPERIMENTAL PROCEDURES

### Animals

D2.*mdx* (Jax# 013141) and D2.WT (Jax# 000671) mice used in this study were originally obtained from Jackson Laboratory. The Nox4^KO^:*mdx* mouse line was created by crossing the *Nox4*^*tm1Kkr*^ knockout allele [(23); Jax# 022996] onto the D2.*mdx* background for 5 generations, as previously reported (15). Following the second backcross, only *Nox4*^*tm1kkr/+*^ mice homozygous for the DBA/2J polymorphism of *Ltbp4* were selected for continued line development (16). This is because both *Nox4* and *Ltbp4* are located on murine chromosome 7, and cross-over is required for both alleles to segregate together. At the completion of the backcrossing, *Nox4* heterozygous breed pairs were mated to produce Nox4^WT^:*mdx* and Nox4^KO^:*mdx* littermates that were used in experiments. All mice were genotyped for *Nox4* using primer sequences provided by Jackson Laboratory, and for *Ltbp4* and *mdx* alleles using published genotyping primers (16).

Mice were housed 3-5 mice per cage, randomly assigned into groups, provided *ad libitum* access to food (NIH-31 Open formulation diet; Envigo #7917), water, and enrichment, and maintained on a 12-hour light/dark system. Once daily (Q.D.) GKT831 (purchased from Ambeed, Inc.) was administered orally (P.O.) suspended at a concentration of 36 mg/mL in a vehicle consisting of sterilized sunflower seed oil (Sigma-Aldrich). This method of drug delivery results in voluntary ingestion of administered solution by mice, similar to syrup-based vehicles as previously reported (17). The GKT831 treatment group received a dose of 60 mg/kg (21). All animal procedures were approved and conducted in accordance with the University of Florida IACUC and reported using the recommendations of ARRIVE guidelines.

### Immunofluorescence and histological evaluations

OCT-embedded murine tissues were cryo-sectioned at 10 µm and fixed in 4% PFA. Immunofluorescent analysis was performed using anti-NOX4 (1:2000; Abcam #13303), anti-laminin (1:800; Novus #MAB2549), anti-vimentin (1:1000; Novus #MB300), anti-PDGFRα (1:500; R&D Systems #AF1062), anti-SMA (1:1000; Abcam #ab5694), anti-periostin (1:500; Abcam #ab14041), anti-Pax7 (1:100; R&D Systems #MAB1675), anti-Myosin X (1:800; Sigma #HPA024223) and anti-active caspase-3 (1:1000; Cell Signaling #9661) primary antibodies. Mouse tissue sections incubated with mouse monoclonal antibodies were first incubated with a solution containing donkey anti-mouse IgG AffiniPure Fab fragments (1:25 in PBS; Jackson ImmunoResearch #715-007-003) for one hour prior to blocking. Following overnight incubation in primary antibody, sections were rinsed with PBS and incubated with the appropriate species-specific secondary antibody (1:500 dilution). Lipofuscin-dependent autofluorescence was quenched using a 0.1% solution of Sudan black B following secondary antibody incubation. Slides were cover-slipped using Prolong Gold mounting reagent (Invitrogen). Images were acquired using a Leica SP8 confocal microscope. All comparative images were stained simultaneously and acquired using identical settings. Human samples were obtained through the National Disease Research Interchange (Philadelphia, PA).

Picrosirius red staining was performed as previously described (14) following decalcification of muscle sections using Formical-2000 (StatLab). Slides were visualized with a Leica DMR microscope, and images were acquired using a Leica DFC310FX camera interfaced with Leica LAS X software. All comparative images were stained simultaneously and acquired using identical settings. Images were processed and analyzed by investigators blinded to study groups using ImageJ software, as previously described (14). Image quantification for each sample consisted of the mean value from five independent and randomly-selected fields of view for each muscle section.

### Assessment of muscle function

Functional assessments of the EDL and diaphragm muscles were evaluated as previously described (17) by the University of Florida Physiological Assessment Core. Muscles of anesthetized mice were dissected and placed in physiological Ringer’s solution gas equilibrated with 95% O_2_ /5% CO_2_. After determining optimum length, muscles were subjected to three isometric contractions (stimulated at 120 Hz for 500 ms) to determine maximum tetanic tension (Po). Following these procedures, muscles were weighed, frozen embedded in OCT or snap-frozen, and stored at −80 °C until further use.

### Collagen Assay

Snap-frozen gastrocnemius samples were pulverized and transferred to pre-weighed 2 mL micro-centrifuge tubes. Tendon pieces were removed from samples during the pulverization procedure. Deionized H_2_O was added to the sample at a volume of 10-times the sample mass, and the tissues were vigorously disrupted using a hand homogenizer (Benchmark Scientific Model D1000). A volume of 200 µL for each resulting sample homogenate was transferred to pre-weighed screw-top micro-centrifuge tubes, and samples were completely desiccated by overnight incubation at 65 °C in order to determine the dry mass of the sample to be assayed. This is important because of potential differences in tissue mass caused by edema that may affect assay results. Following tissue dry mass determination, the collagen content of the samples was measured using a colorimetric Total Collagen Assay kit (Biovision #K218), following the manufacturer’s directions in a 96-well plate. Assay data were collected using a SpectraMax i3x multi-mode spectrophotometer (Molecular Devices) at a wavelength of 560 nm.

### Gene Expression

Gene expression analysis was conducted as previously described (17) using the following mouse-specific primers: *Col1a2* (forward) 5’-ATG GTG GCA GCC AGT TTG AA-3’ and (reverse) 5’-TCC AGG TAC GCA ATG CTG TT-3’; *Acta2* (forward) 5’-CTA CTG CCG AGC GTG AGA TTG TCC-3’ and (reverse) 5’-GAG GGC CCA GCT TCG TCG TAT T-3’; *Il6* (forward) 5’-AAC CAC GGC CTT CCC TAC TTC-3’ and (reverse) 5’-TCT GGC TTT GTC TTT CTT GTT ATC-3’; *Tgfb1* (forward) 5’-GAC TCT CCA CCT GCA AGA CCA T-3’ and (reverse) 5’-GGG ACT GGC GAG CCT TAG TT; *Gapdh* (forward) 5’-AGC AGG CAT CTG AGG GCC CA-3’ and (reverse) 5’-TGT TGG GGG CCG AGT TGG GA-3’. Relative gene expression quantification was performed using the ΔΔCt method with *Gapdh* as the normalization gene.

### Immunoblotting

Immunoblotting was performed as previously described (17) using the following primary antibodies: anti-dystrophin (1:1000; Abcam #15277), anti-NOX4 (1:2000; Abcam #13303), anti-fibronectin (1:2000; Sigma #F7387), and anti-GAPDH (1:2000; Santa Cruz #sc-25778). Quantified band signal intensities were measured using Image Studio Lite software (LI-COR Biosciences), normalized to GAPDH signal values, and reported relative to respective control samples.

### Statistical Analysis

Statistical analysis was performed using unpaired, two-tailed Welch’s T-test [α = 0.05; effect size reported as Cohen’s *d* (*d*)] or ANOVA (Tukey post-hoc tests; α = 0.05; effect size reports as η^2^). A P value less than 0.05 was considered significant. Values are displayed as box-and-whisker plots (depicting minimum and maximum values) or as bar graphs showing mean ± SEM.

## ACKNOWLEDGEMENTS

This work was funded by a grant from the Muscular Dystrophy Association (MDA549004) to DWH. Michael Matheny, Lillian Wright, Megan Armellini, and Shailja Desai are thanked for their technical assistance during this work.

## CONFLICT OF INTEREST

The author declares that there are no conflicts of interest with the contents of this article.

## SUPPLEMENTAL FIGURES

**Figure S1.**
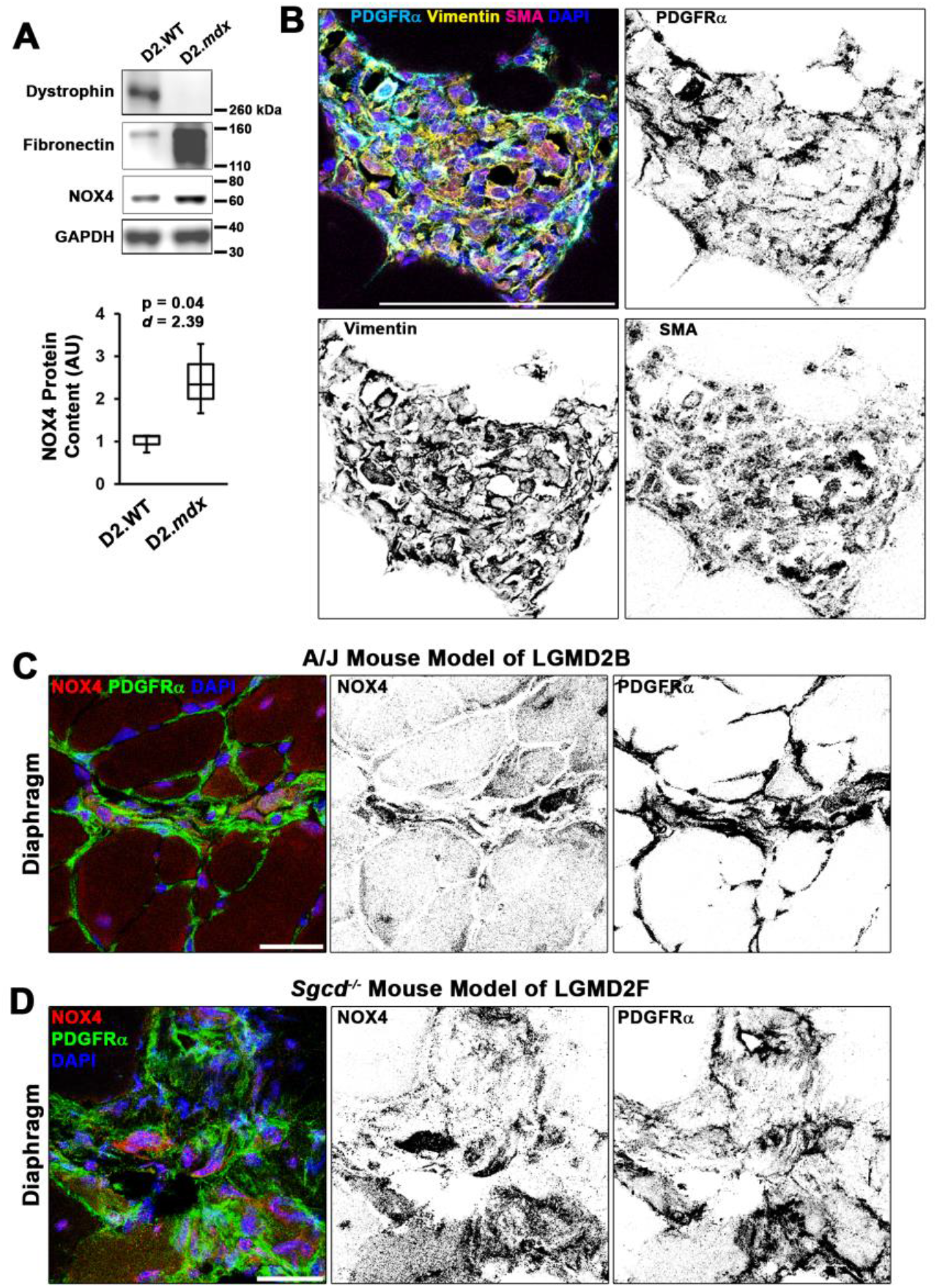
NOX4 expression in dystrophic muscle. Related to Figure 1. (**A**) Representative immunoblots display differential content of dystrophin, fibronectin, and NOX4 protein levels in quadriceps lysates from 4 mo male D2.WT and D2.*mdx* mice (n = 4). NOX4 immunoblotting data were quantified, normalized to GAPDH signal, and displayed relative to D2.WT values. Data are presented as box-and-whisker plots, with error bars representing minimum and maximum values, and were analyzed using unpaired, two-tailed Welch’s T-tests (α = 0.05; effect size is presented as Cohen’s *d*). (**B**) Immunofluorescent detection of myofibroblasts in D2.*mdx* gastrocnemius muscles using antibodies against PDGFRα, vimentin, and SMA. NOX4 localizes to PDGFRα^+^ cells in other models of muscular dystrophies, including diaphragms of (**C**) 9 month-old A/J mice, a model of LGMD2B and (**D**) 3 month-old *Sgcd*^*-/-*^ mice, a model of LGMD2F. Scale bars represent 50 µm.

**Figure S2.**
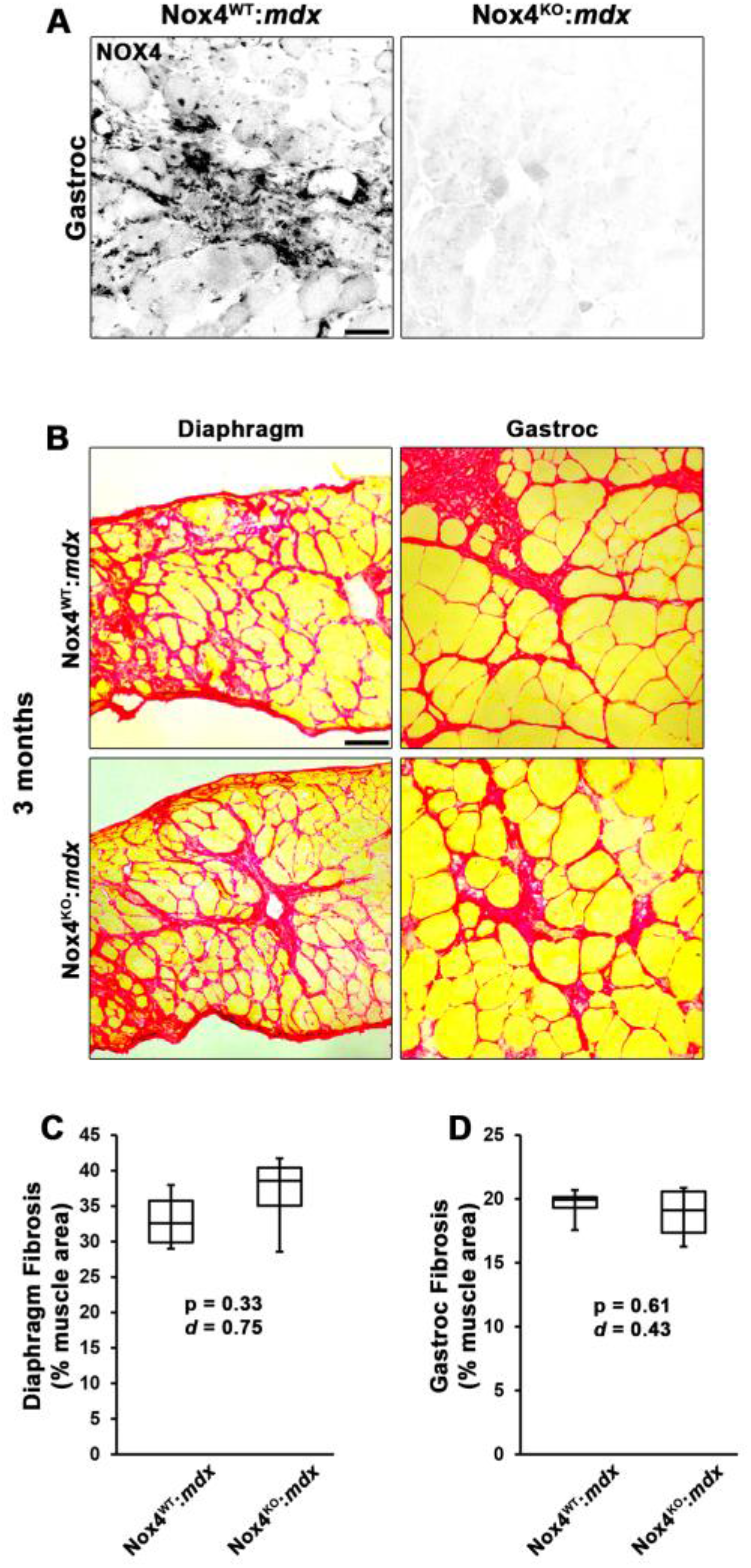
NOX4 ablation does not affect early time course of muscle fibrosis. Related to Figure 2. (**A**) Representative immunofluorescent images from Nox4^WT^:*mdx* and Nox4^KO^:*mdx* gastrocnemius (gastroc) muscles stained with anti-NOX4 antibody. Muscle fibrosis was histologically assessed in the diaphragm and gastroc muscles from 3 month-old Nox4^WT^:*mdx* and Nox4^KO^:*mdx* mice using picrosirius red staining (n = 4). (**B**) Representative images and (**C-D**) histological quantifications reveal no significant differences in muscle fibrosis between the groups at this age. Data are presented as box-and-whisker plots with error bars representing minimum and maximum values. Data were analyzed using unpaired, two-tailed Welch’s T-tests (α = 0.05; effect size is presented as Cohen’s *d*). Scale bar represents 100 µm.

## Notes

### Competing Interest Statement

The authors have declared no competing interest.

## REFERENCES

1. Mendell JR, et al. (2012) Evidence-based path to newborn screening for Duchenne muscular dystrophy. Ann Neurol 71(3):304–313.

2. Hoffman EP, Brown RH, Jr., & Kunkel LM (1987) Dystrophin: the protein product of the Duchenne muscular dystrophy locus. Cell 51(6):919–928.

3. Petrof BJ, Shrager JB, Stedman HH, Kelly AM, & Sweeney HL (1993) Dystrophin protects the sarcolemma from stresses developed during muscle contraction. Proc Natl Acad Sci U S A 90(8):3710–3714.

4. Willcocks RJ, et al. (2016) Multicenter prospective longitudinal study of magnetic resonance biomarkers in a large duchenne muscular dystrophy cohort. Ann Neurol 79(4):535–547.

5. Charge SB & Rudnicki MA (2004) Cellular and molecular regulation of muscle regeneration. Physiol Rev 84(1):209–238.

6. Mauro A (1961) Satellite cell of skeletal muscle fibers. J Biophys Biochem Cytol 9:493–495.

7. Arnold L, et al. (2007) Inflammatory monocytes recruited after skeletal muscle injury switch into antiinflammatory macrophages to support myogenesis. J Exp Med 204(5):1057–1069.

8. Fry CS, Kirby TJ, Kosmac K, McCarthy JJ, & Peterson CA (2017) Myogenic Progenitor Cells Control Extracellular Matrix Production by Fibroblasts during Skeletal Muscle Hypertrophy. Cell Stem Cell 20(1):56–69.

9. Wosczyna MN, et al. (2019) Mesenchymal Stromal Cells Are Required for Regeneration and Homeostatic Maintenance of Skeletal Muscle. Cell Rep 27(7):2029–2035 e2025.

10. Hardy D, et al. (2016) Comparative Study of Injury Models for Studying Muscle Regeneration in Mice. PLoS One 11(1):e0147198.

11. McCarthy JJ, et al. (2011) Effective fiber hypertrophy in satellite cell-depleted skeletal muscle. Development 138(17):3657–3666.

12. Patsalos A, et al. (2017) In situ macrophage phenotypic transition is affected by altered cellular composition prior to acute sterile muscle injury. J Physiol 595(17):5815–5842.

13. Dadgar S, et al. (2014) Asynchronous remodeling is a driver of failed regeneration in Duchenne muscular dystrophy. J Cell Biol 207(1):139–158.

14. Hammers DW, et al. (2020) The D2.mdx mouse as a preclinical model of the skeletal muscle pathology associated with Duchenne muscular dystrophy. Sci Rep 10(1):14070.

15. Fukada S, et al. (2010) Genetic background affects properties of satellite cells and mdx phenotypes. Am J Pathol 176(5):2414–2424.

16. Coley WD, et al. (2016) Effect of genetic background on the dystrophic phenotype in mdx mice. Hum Mol Genet 25(1):130–145.

17. Hammers DW, et al. (2020) Glucocorticoids counteract hypertrophic effects of myostatin inhibition in dystrophic muscle. JCI Insight 5(1).

18. Bernard K & Thannickal VJ (2020) NADPH Oxidase Inhibition in Fibrotic Pathologies. Antioxidants & Redox Signaling 33(6):455–479.

19. Spurney CF, et al. (2008) Dystrophin-deficient cardiomyopathy in mouse: expression of Nox4 and Lox are associated with fibrosis and altered functional parameters in the heart. Neuromuscul Disord 18(5):371–381.

20. Hinz B & Lagares D (2019) Evasion of apoptosis by myofibroblasts: a hallmark of fibrotic diseases. Nature Reviews Rheumatology 16(1):11–31.

21. Gorin Y, et al. (2015) Targeting NADPH oxidase with a novel dual Nox1/Nox4 inhibitor attenuates renal pathology in type 1 diabetes. Am J Physiol Renal Physiol 308(11):F1276–1287.

22. Jha JC, et al. (2014) Genetic targeting or pharmacologic inhibition of NADPH oxidase nox4 provides renoprotection in long-term diabetic nephropathy. J Am Soc Nephrol 25(6):1237–1254.

23. Carnesecchi S, et al. (2011) A key role for NOX4 in epithelial cell death during development of lung fibrosis. Antioxid Redox Signal 15(3):607–619.

24. Hecker L, et al. (2009) NADPH oxidase-4 mediates myofibroblast activation and fibrogenic responses to lung injury. Nat Med 15(9):1077–1081.

25. Jarman ER, et al. (2014) An inhibitor of NADPH oxidase-4 attenuates established pulmonary fibrosis in a rodent disease model. Am J Respir Cell Mol Biol 50(1):158–169.

26. Jiang JX, et al. (2012) Liver fibrosis and hepatocyte apoptosis are attenuated by GKT137831, a novel NOX4/NOX1 inhibitor in vivo. Free Radic Biol Med 53(2):289–296.

27. Lan T, Kisseleva T, & Brenner DA (2015) Deficiency of NOX1 or NOX4 Prevents Liver Inflammation and Fibrosis in Mice through Inhibition of Hepatic Stellate Cell Activation. PLoS One 10(7):e0129743.

28. Somanna NK, et al. (2016) The Nox1/4 Dual Inhibitor GKT137831 or Nox4 Knockdown Inhibits Angiotensin-II-Induced Adult Mouse Cardiac Fibroblast Proliferation and Migration. AT1 Physically Associates With Nox4. J Cell Physiol 231(5):1130–1141.

29. Reutens AT, et al. (2020) A physician-initiated double-blind, randomised, placebo-controlled, phase 2 study evaluating the efficacy and safety of inhibition of NADPH oxidase with the first-in-class Nox-1/4 inhibitor, GKT137831, in adults with type 1 diabetes and persistently elevated urinary albumin excretion: Protocol and statistical considerations. Contemp Clin Trials 90:105892.

30. Joe AW, et al. (2010) Muscle injury activates resident fibro/adipogenic progenitors that facilitate myogenesis. Nat Cell Biol 12(2):153–163.

31. Uezumi A, Fukada S, Yamamoto N, Takeda S, & Tsuchida K (2010) Mesenchymal progenitors distinct from satellite cells contribute to ectopic fat cell formation in skeletal muscle. Nat Cell Biol 12(2):143–152.

32. Contreras O, et al. (2019) Cross-talk between TGF-beta and PDGFRalpha signaling pathways regulates the fate of stromal fibro-adipogenic progenitors. J Cell Sci 132(19).

33. Li R, et al. (2018) Pdgfra marks a cellular lineage with distinct contributions to myofibroblasts in lung maturation and injury response. Elife 7:e36865.

34. Hu B & Phan SH (2013) Myofibroblasts. Curr Opin Rheumatol 25(1):71–77.

35. Aoyama T, et al. (2012) Nicotinamide adenine dinucleotide phosphate oxidase in experimental liver fibrosis: GKT137831 as a novel potential therapeutic agent. Hepatology 56(6):2316–2327.

36. Kanisicak O, et al. (2016) Genetic lineage tracing defines myofibroblast origin and function in the injured heart. Nat Commun 7:12260.

37. Hara M, et al. (2018) Periostin Promotes Fibroblast Migration and Inhibits Muscle Repair After Skeletal Muscle Injury. J Bone Joint Surg Am 100(16):e108.

38. Lorts A, Schwanekamp JA, Baudino TA, McNally EM, & Molkentin JD (2012) Deletion of periostin reduces muscular dystrophy and fibrosis in mice by modulating the transforming growth factor-beta pathway. Proc Natl Acad Sci U S A 109(27):10978–10983.

39. Thooyamani AS & Mukhopadhyay A (2021) PDGFRalpha mediated survival of myofibroblasts inhibit satellite cell proliferation during aberrant regeneration of lacerated skeletal muscle. Sci Rep 11(1):63.

40. Kottlors M & Kirschner J (2010) Elevated satellite cell number in Duchenne muscular dystrophy. Cell Tissue Res 340(3):541–548.

41. He WA, et al. (2013) NF-kappaB-mediated Pax7 dysregulation in the muscle microenvironment promotes cancer cachexia. J Clin Invest 123(11):4821–4835.

42. Hammers DW, et al. (2021) Filopodia powered by class x myosin promote fusion of mammalian myoblasts. Elife 10:e72419.

43. Hecker L, et al. (2014) Reversal of persistent fibrosis in aging by targeting Nox4-Nrf2 redox imbalance. Sci Transl Med 6(231):231ra247.

44. Allen RE & Boxhorn LK (1987) Inhibition of skeletal muscle satellite cell differentiation by transforming growth factor-beta. J Cell Physiol 133(3):567–572.

45. Mazala DA, et al. (2020) TGF-beta-driven muscle degeneration and failed regeneration underlie disease onset in a DMD mouse model. JCI Insight 5(6):e135703.

46. Duan D (2018) Systemic AAV Micro-dystrophin Gene Therapy for Duchenne Muscular Dystrophy. Mol Ther 26(10):2337–2356.

47. England SB, et al. (1990) Very mild muscular dystrophy associated with the deletion of 46% of dystrophin. Nature 343(6254):180–182.

48. Barp A, et al. (2017) Muscle MRI and functional outcome measures in Becker muscular dystrophy. Sci Rep 7(1):16060.

